# Identification of a discrete neuronal circuit that relays insulin signaling into the brain to regulate glucose homeostasis

**DOI:** 10.1101/2021.09.13.460055

**Authors:** Mouna El Mehdi, Saloua Takhlidjt, Mélodie Devère, Arnaud Arabo, Marie-Anne Le Solliec, Julie Maucotel, Alexandre Bénani, Emmanuelle Nedelec, Céline Duparc, Benjamin Lefranc, Jérôme Leprince, Youssef Anouar, Gaëtan Prévost, Nicolas Chartrel, Marie Picot

**Author notes:** **Corresponding author**: Dr Marie Picot, Normandie Univ, UNIROUEN, INSERM U1239, Laboratory of Neuronal and Neuroendocrine Differentiation and Communication (DC2N), 76000 Rouen, France.

## Abstract

26RFa (QRFP) is a biologically active peptide that regulates glucose homeostasis by acting as an incretin and by increasing insulin sensitivity at the periphery. 26RFa is also produced by a neuronal population localized in the hypothalamus. In the present study, we have investigated whether the 26RFa neurons may be involved in the hypothalamic regulation of glucose homeostasis. Our data indicate that 26RFa, i.c.v. injected, induces a robust antihyperglycemic effect associated with an increase of insulin production by the pancreatic islets. In addition, we found that insulin strongly stimulates 26RFa expression and secretion by the hypothalamus. RNAscope experiments revealed that neurons expressing 26RFa in the lateral hypothalamic area and the ventromedial hypothalamic nucleus also express the insulin receptor and that insulin induces the expression of 26RFa in these neurons. Concurrently, we show that the central antihyperglycemic effect of insulin is abolished in presence of a 26RFa receptor (GPR103) antagonist as well as in mice deficient for 26RFa. Finally, our data indicate that the hypothalamic 26RFa neurons are not involved in the central inhibitory effect of insulin on hepatic glucose production, but mediate the central effects of the hormone on its own peripheral production.

To conclude, in the present study we have identified a novel actor of the hypothalamic regulation of glucose homeostasis, the 26RFa/GPR103 system and we provide the evidence that this neuronal peptidergic system is a key relay for the central regulation of glucose metabolism by insulin.

## Introduction

Since the discovery of insulin in 1921, research on the regulation of glucose homeostasis and diabetes mellitus has focused on the concept that function of the pancreatic islets is both necessary and sufficient to explain glucose homeostasis, and that diabetes results from defects of insulin secretion, action or both. However, it is known since the 19th century that the brain, and more specifically the hypothalamus, is involved in the control of glucose homeostasis and diabetes pathogenesis (1, 2). It is even suspected that the brain accounts for 50% of overall glucose disposal and can normalize glycemia in rodent models of diabetes (3, 4). From these findings, emerged a new model in which the control of glucose homeostasis results from a complex and highly coordinated interactions between the brain and the pancreatic islets. The crucial role of the brain in the control of glucose metabolism is supported by the observations that injection of glucose, leptin or insulin into discrete hypothalamic areas including the arcuate nucleus (Arc), the ventromedial hypothalamic nucleus (VMH) and the paraventricular nucleus (PVN) lowers blood glucose levels, increases liver insulin sensitivity (2, 5, 6), and that central insulin decreases hepatic glucose production (HGP) (7). Similar effects are achieved by restoring functional insulin receptors to specific hypothalamic nuclei that otherwise lack them (8). Conversely, deletion of insulin receptors or their downstream signalling intermediates from defined hypothalamic neurons induce glucose intolerance and systemic insulin resistance (9, 10). More specifically, it was found that the central effects of insulin on glucose homeostasis are mediated only in part by the AgRP neurons of the Arc and not by the POMC neurons (10, 11). These observations suggest that insulin targets other neuropeptidergic systems in the hypothalamus to exert its central action on glucose homeostasis and highlight the fact that our knowledge on the molecular identity of the hypothalamic neuronal populations relaying insulin central action remains fragmentary.

In this context, the neuropeptidergic 26RFa/GPR103 system, is of particular interest. 26RFa (also referred to as QRFP) is a hypothalamic neuropeptide discovered concurrently by us and others (12-14), and identified as the cognate ligand of the human orphan G protein-coupled receptor GPR103 (13-17). Neuroanatomical observations revealed that 26RFa- and GPR103-expressing neurons are primarily localized in hypothalamic nuclei involved in the control of feeding behaviour including the VMH, the lateral hypothalamic area (LH) and the Arc (12, 16, 18, 19). Indeed, i.c.v. administration of 26RFa stimulates food intake (12, 16, 20, 21), and the neuropeptide exerts its orexigenic activity by modulating the NPY/POMC system in the Arc (21). More recently, the involvement of the 26RFa/GPR103 neuropeptidergic system in the control of glucose homeostasis at the periphery was reported. We, and others, found that 26RFa and GPR103 are strongly expressed by β cells of the pancreatic islets (22-24), and that the neuropeptide prevents cell death and apoptosis of β cells (22). We also showed that 26RFa is abundantly expressed all along the gut and that i.p. administration of the neuropeptide attenuates glucose-induced hyperglycemia by increasing plasma insulin via a direct insulinotropic effect on the pancreatic β cells, and by increasing insulin sensitivity (23, 24). Finally, we reported that an oral glucose challenge induces a massive secretion of 26RFa by the gut into the blood, strongly suggesting that this neuropeptide regulates glycemia by acting as an incretin (23). This incretin effect of 26RFa has been very recently confirmed by the observation that administration of a GPR103 antagonist reduces the global glucose-induced incretin effect, and also decreases insulin sensitivity (25), and that 26RFa mutant mice show an impaired regulation of glucose homeostasis (26).

Altogether, these observations prompted us to investigate whether the hypothalamic 26RFa/GPR103 system is involved in the central regulation of glucose homeostasis and may be a target for insulin.

## Results

### Effect of central administration of 26RFa on glucose homeostasis

After a 6-h or a 12-h fasting, i.c.v. treatment with 26RFa did not affect basal plasma glucose levels during the 90 min period of the test (Fig. 1A, B). By contrast, a glucose tolerance test (IPGTT) revealed that central administration of 26RFa significantly attenuates (p<0.01) the hyperglycemia induced by an i.p. glucose challenge all along the duration of the test (Fig. 1C). Administration of the 26RFa receptor antagonist 25e, strongly diminished the antihyperglycemic effect of 26RFa, whereas 25e alone had no effect on plasma glucose levels (Fig. 1D). Concurrently, an acute glucose-stimulated insulin secretion test revealed that i.c.v. injection of 26RFa significantly potentiated (p<0.01) glucose-induced insulin production (Fig. 1E). Finally, 26RFa had no effect on insulin-induced hypoglycemia during the first 30 min of an insulin tolerance test, but the peptide attenuated (p<0.05) the hypoglycemic action of insulin during the last phase of the test (Fig. 1F).

**Figure 1.**
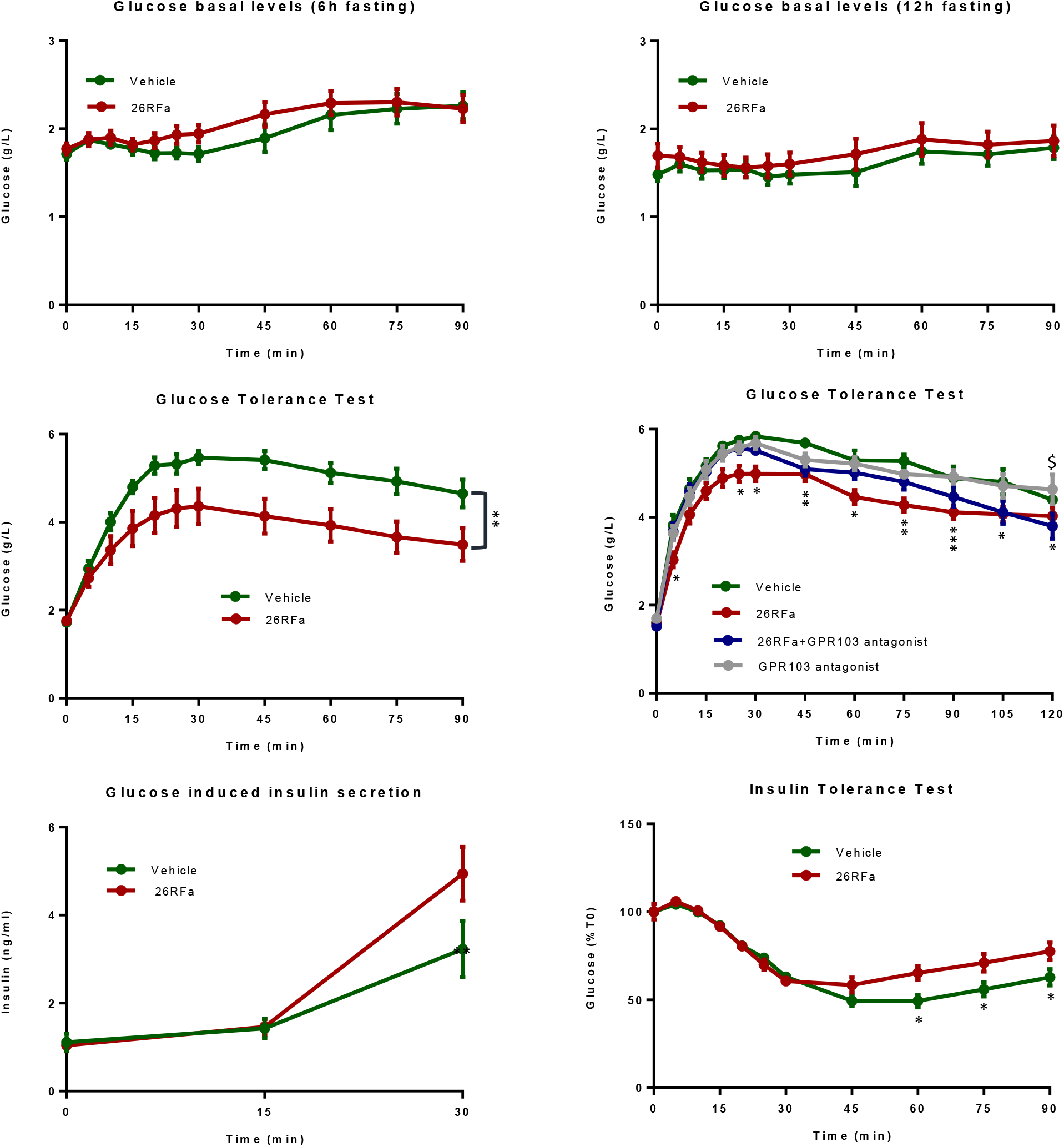
Effect of central administration of 26RFa on glucose homeostasis. **A, B:** Effect of i.c.v. administration of 26RFa (3 µg) on plasma glucose levels in basal condition after a 6-h or 12-h fasting period (n=10-15). **C, D:** Effect of i.c.v. administration of 26RFa alone (3 µg) (C) or together with the GPR103 antagonist 25e (10^−4^ M) (D) on plasma glucose levels during an IPGTT (n=15). **E:** Effect of i.c.v. administration of 26RFa (3 µg) on insulin production during an AGSIS (n=11). **F:** Effect of i.c.v. administration of 26RFa (3 µg) on plasma glucose levels during an ITT (n=16). Data represent means ± SEM of 5 independent experiments for each test. *, p<0.05; **, p<0.01; *** p<0.001 (26RFa *vs* vehicle).

### Effect of glucose, leptin and insulin on 26RFa-producing neurons

The impact of factors well known to regulate glucose homeostasis such as glucose, leptin and insulin on hypothalamic 26RFa-producing neurons was investigated using *in vitro* and *ex vivo* complementary approaches. Firstly, we used the hypothalamic neuronal cell line m-HypoA-59 that expresses 26RFa and its receptor GPR103 as a model of 26RFa-producing neurons (27-29). Our data revealed that glucose had no effect on the secretion of 26RFa and GPR103 expression by these cells (Fig. 2A, B). Leptin increased significantly 26RFa secretion (p<0.05) by the m-HypoA-59 cells at a dose of 100 nM, but did not alter the expression of GPR103 (Fig. 2C, D). Insulin enhanced significantly (p<0.05) 26RFa secretion by the cells at a dose of 10 nM but did not impact the expression of the 26RFa receptor (Fig. 2E, F). The same parameters were examined using hypothalamic explants. As observed on the m-HypoA-59 cell line, we found that glucose does not significantly alter the secretion and expression of 26RFa and its receptor by the whole hypothalamus (Fig. 2G-I). Leptin did not alter the secretion of the 26RFa by the hypothalamic explants (Fig. 2J) but stimulates significantly (p<0.05) the expression of the peptide at a dose of 10 nM (Fig. 2K). The expression of GPR103 was not altered by leptin (Fig. 2L). Finally, we found that insulin strongly stimulated dose-depently (p<0.01) 26RFa release by the hypothalamic explants (Fig. 2M) without modifying the expression of the peptide or its receptor (Fig. 2N, O).

**Figure 2.**
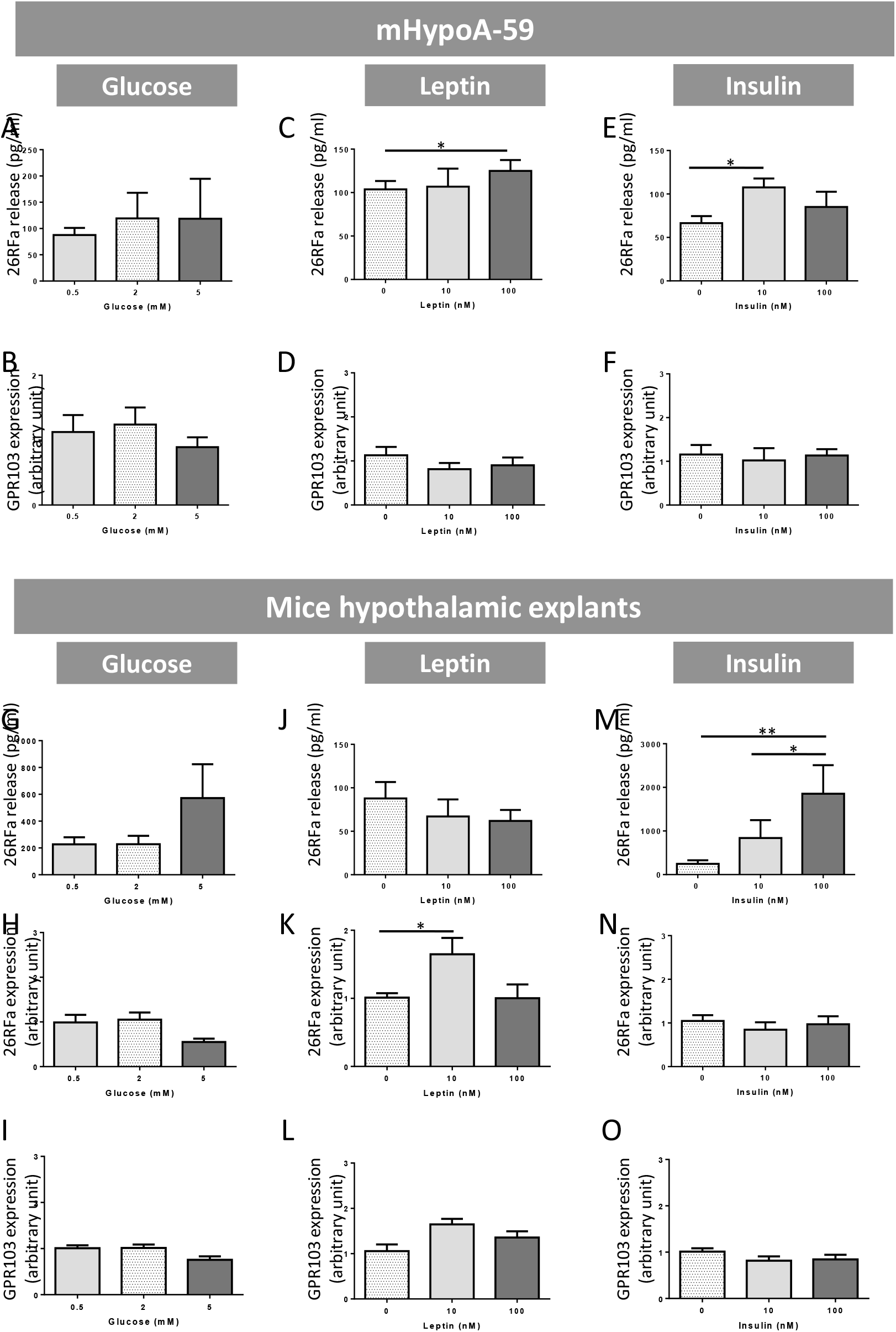
Effect of glucose, leptin and insulin on 26RFa-producing neurons. **A-F:** Effect of various concentrations of glucose, leptin and insulin on 26RFa secretion and GPR103 expression by the hypothalamic neuronal cell line m-HypoA-59. **G-O:** Effect of various concentrations of glucose, leptin and insulin on the secretion and expression of 26RFa, and the expression of GPR103 by hypothalamic explants. Data represent means ± SEM of 4 independent experiments (n=9/condition). *, p<0.05; **, p<0.01compared to control conditions (grey bars).

### Neuroanatomical interactions between insulin and 26RFa-expressing neurons

The neuroanatomical interactions between insulin and 26RFa-expressing neurons were investigated using RNAscope® technology. Figure 3A shows a panoramic view of the hypothalamus at the level of the Arc as assessed by the strong AgRP (green labelling) that revealed the presence of scattered 26RFa-expressing neurons (red labelling) restricted to the VMH and LH, whereas the InsR (white labelling) is widely distributed throughout the slice. At a higher magnification, it appeared that the Arc exhibited a strong AgRP (as expected) and InsR labelling but was totally devoid of 26RFa-expressing neurons (Fig. 3B). By contrast, a number of 26RFa-expressing neurons were detected in the VMH and the LH with a majority of these neurons showing a low expression of the neuropeptide, although some of them were strongly labelled (Fig. 3C, D). As already observed in the Arc, the VMH and the LH exhibited a strong expression of the InsR (Fig. 3C, D). In addition, selected photomicrographs show a co-expression of the InsR and 26RFa in neurons of the VMH (Fig. 4A) and the LH (Fig. 4B). In a second step, we investigated the impact of i.c.v. administration of insulin during an IPGTT on the expression of 26RFa, GPR103 and InsR using the qRT-PCR and RNAscope approaches. The qRT-PCR experiment revealed that central injection of insulin increased significantly (p<0.05) the hypothalamic expression of 26RFa, but had no effect on GPR103 and InsR mRNA levels (Fig. 5A). Representative panoramic views taken during the RNAscope procedure in the anterior part of the hypothalamus confirmed the qRT-PCR data and showed the occurrence of strongly labelled 26RFa-expressing neurons in the LH and VMH of brains receiving insulin (Fig. 5B, C). Focus on the LH and the VMH revealed the increase in the number of 26RFa-expressing neurons and/or the level of expression the neuropeptide in these neurons (Fig. 6A, B). Interestingly, we also found that insulin induced the occurrence of 26RFa-expressing neurons in the Arc (Fig. 6C).

**Figure 3.**
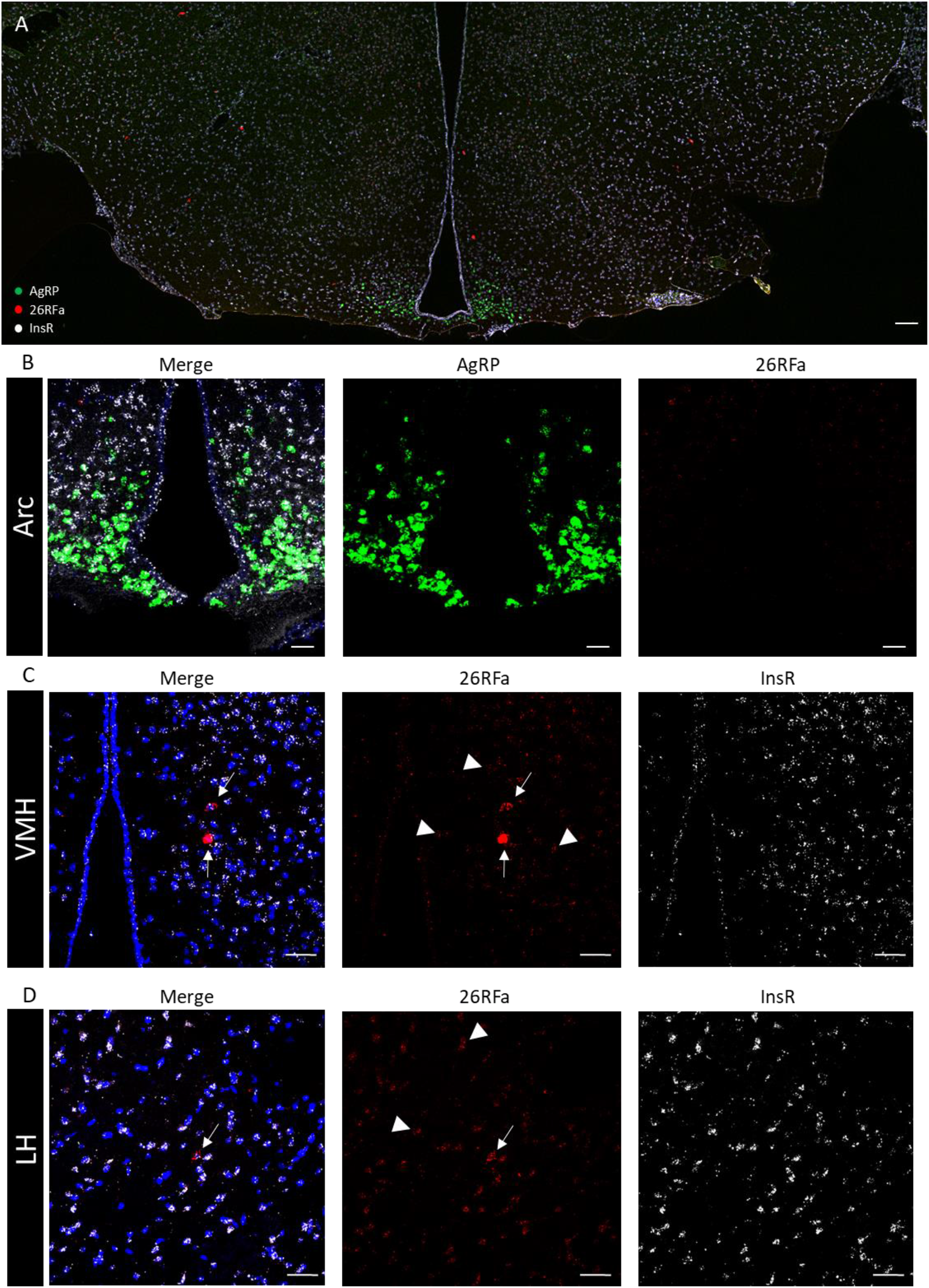
Distribution of 26RFa-, AgRP- and InsR-expressing neurons in the hypothalamus by using the RNAscope approach. **A:** Panoramic view of a hypothalamic section (at the level of the arcuate nucleus (Arc)) showing the presence of 26RFa-expressing neurons (red spots) in the ventromedial hypothalamic nucleus (VMH) and the lateral hypothalamic area (LH). AgRP-expressing neurons (green spots) are strictly confined to the Arc whereas insulin receptor (InsR) mRNA (white spots) are widely distributed throughout the whole hypothalamus. **B:** Focus on the Arc reveals a robust AgRP and InsR labelling and the absence of 26RFa-expressing neurons. **C, D:** Focus on the VMH (C) and the LH (D) reveals the presence of numerous 26RFa-expressing neurons, most of them exhibiting a low expression of the neuropeptide (arrowheads) but a few showing a strong labelling (arrows). The InsR is widely distributed throughout the VMH and the LH. Scale bars: (A)=200 µm; (B-D)=25 µm.

**Figure 4.**
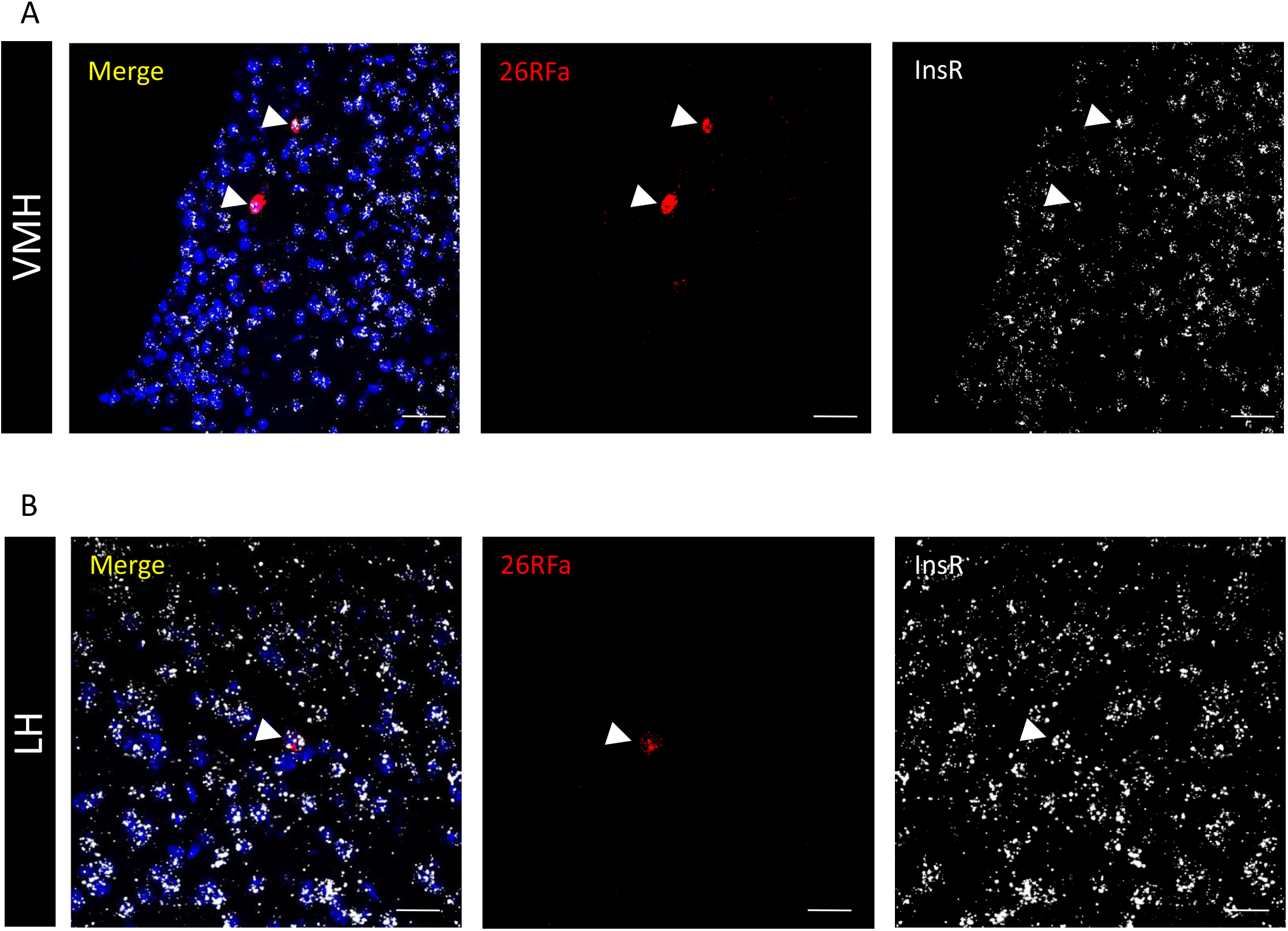
Co-expression of 26RFa with the InsR in the hypothalamus. **A, B:** Representative photomicrographs of double labelling in the ventromedial hypothalamic nucleus (VMH) (A) and the lateral hypothalamic area (LH) (B) showing that 26RFa (red spots) and the insulin receptor (InsR, white spots) are co-expressed in the same neurons (arrows). Scale bars: 25 µm.

**Figure 5.**
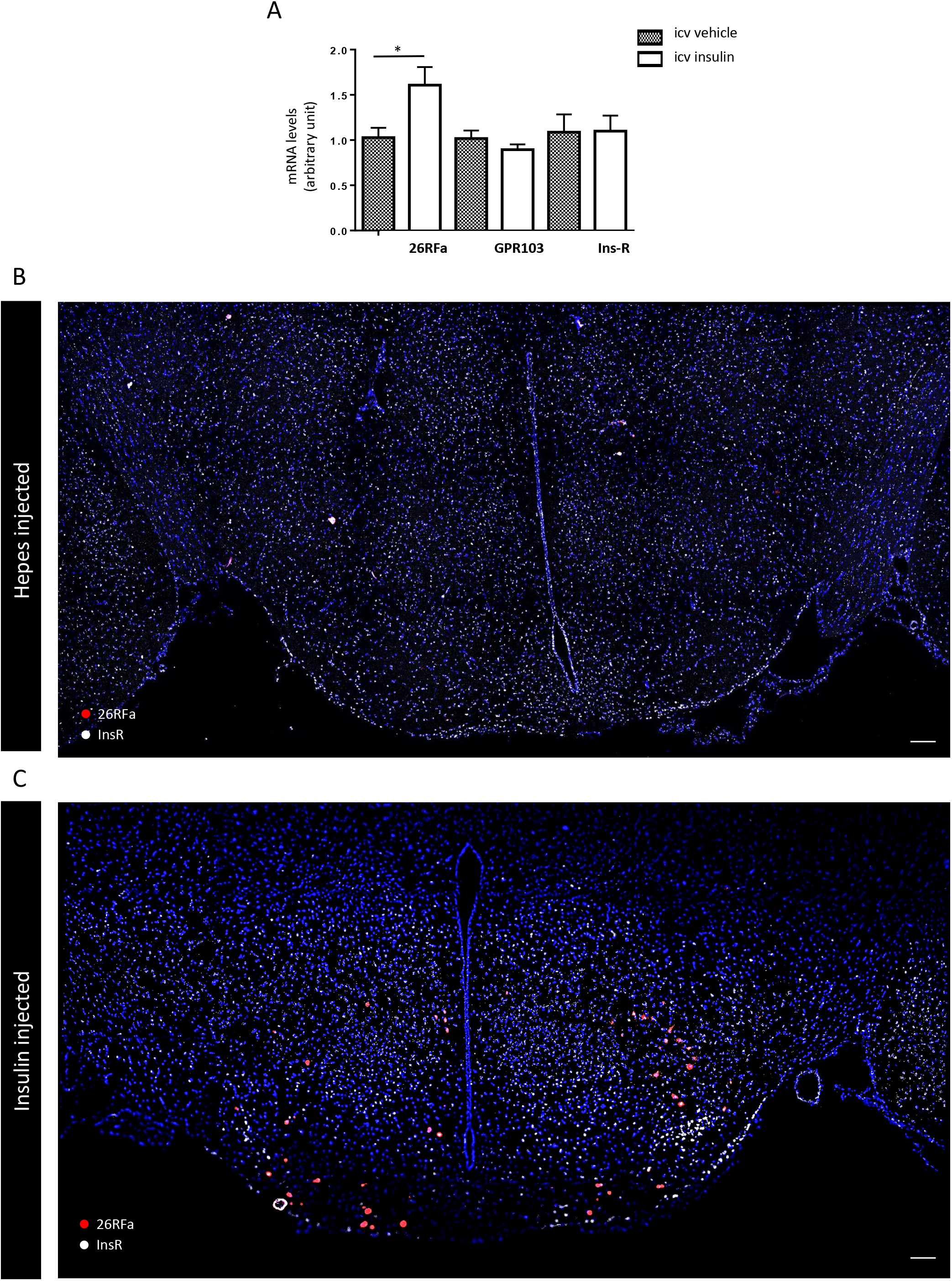
Effect of i.c.v. administration of insulin on the expression of 26RFa in the hypothalamus. **A:** Effect of i.c.v. administration of insulin (10 mUI) on the hypothalamic expression of 26RFa, GPR103 and the insulin receptor (InsR) showing an important increase of 26RFa mRNA levels and no alteration of GPR103 and InsR mRNA levels. Data represent means ± SEM of 6 hypothalami per condition. *, p<0.05. **B, C:** Panoramic views of sections through the anterior part of the hypothalamus of brains injected with hepes (B) or insulin (C) showing the occurrence of strongly labelled 26RFa-expressing neurons (red spots) in the LH and VMH of insulin-treated brains. Scale bars: 200 µm.

**Figure 6.**
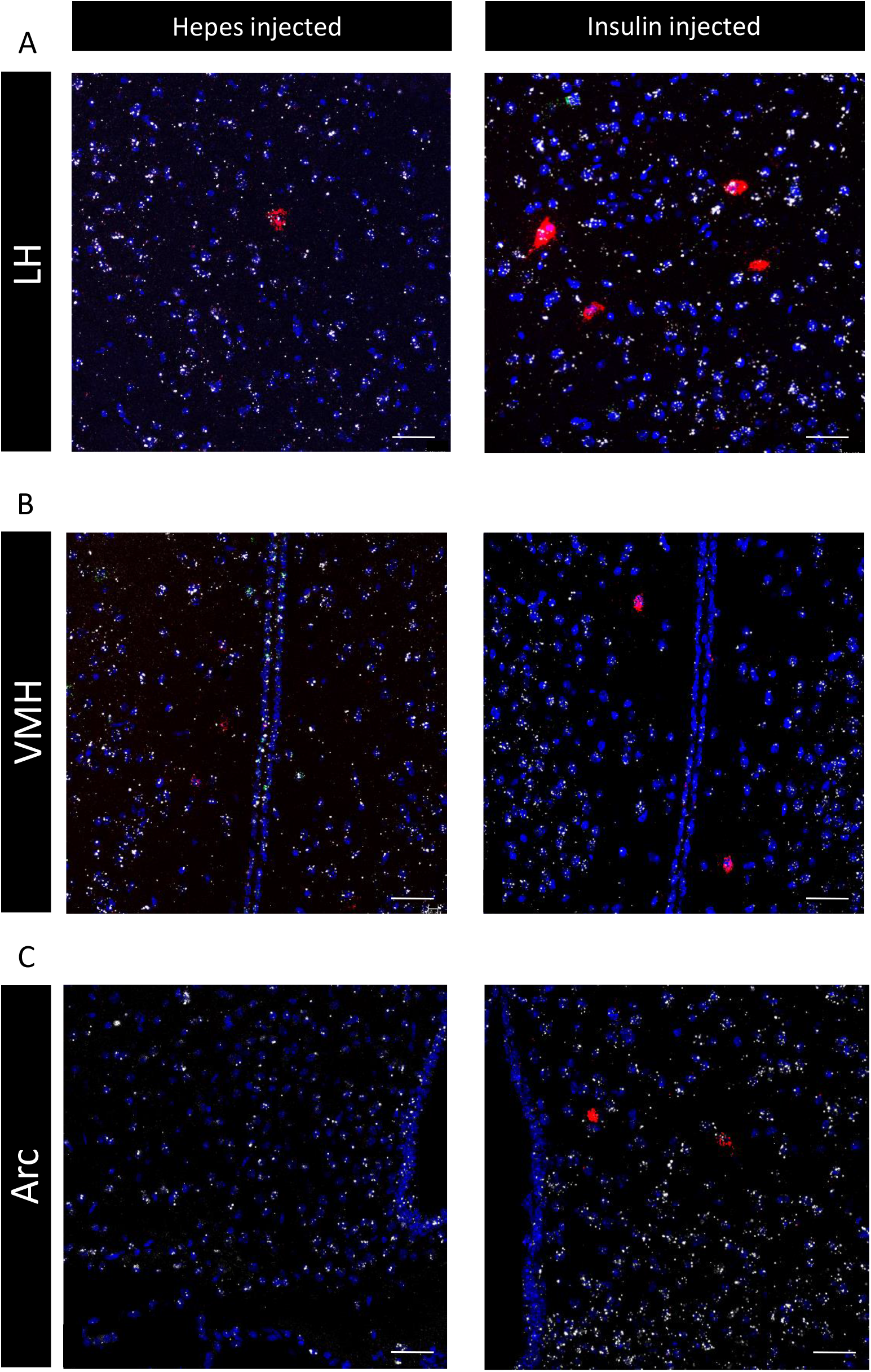
Effect of i.c.v. administration of insulin on the expression of 26RFa in various hypothalamic nuclei. **A, B:** Representative photomicrographs of the lateral hypothalamic area (LH) (A) and the ventromedial hypothalamic nucleus (VMH) (B) of hepes- or insulin-injected brains showing the increase in the number of 26RFa-expressing neurons (red spots) and/or the level of expression the neuropeptide induced by insulin (10 mUI). The white spots correspond the insulin receptor (InsR) mRNAs. **C:** Representative photomicrographs showing that insulin induces the occurrence of 26RFa-expressing neurons in the Arc. Scale bars: 25 µm.

### Effect of blocking the 26RFa/GPR103 system on insulin hypothalamic signaling

An IPGTT revealed that i.c.v. administration of insulin induced a robust significant antihyperglycemic effect (p<0.05 to p<0.001) (Fig. 7A). Interestingly, 26RFa displayed an antihyperglycemic profile very similar in terms of amplitude and kinetic, to that of insulin (Fig. 7A). Co-administration of the GPR103 antagonist 25e with insulin during an IPGTT resulted in a total blockade of the antihyperglycemic effect of insulin during the first 60 min of the test whereas the administration of 25e alone did not affect the glycemic profile (Fig. 7B). In addition, an IPGTT performed in mice deficient for 26RFa revealed a loss of the antihyperglycemic effect of insulin, as compared to the injection of the hormone in wild type animals (Fig. 7C).

**Figure 7.**
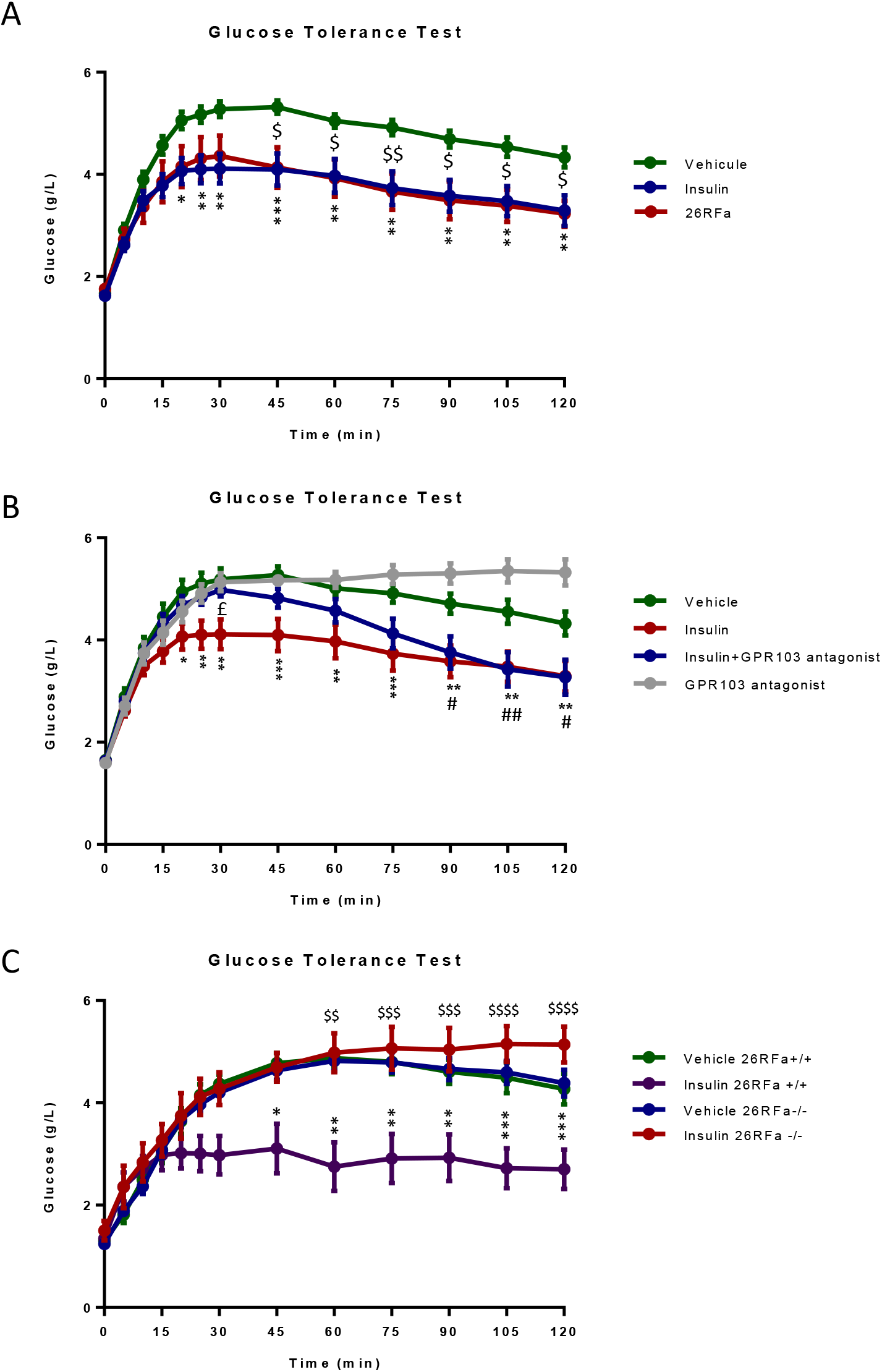
Central interaction between insulin and the hypothalamic 26RFa/GPR103 system during a glucose tolerance test. **A:** Effect of i.c.v. administration of insulin (10 mUI) or 26RFa (3 µg) on plasma glucose levels during a glucose tolerance test (IPGTT). **B:** Effect of i.c.v. administration of insulin (10 mUI) alone, or with the GPR103 receptor antagonist 25e (10^−4^ M) on plasma glucose levels during an IPGTT. **C:** Effect of i.c.v. administration of insulin (10 mUI) on plasma glucose levels during an IPGTT in 26RFa ^+/+^ and 26RFa^-/-^ mice. Data represent means ± SEM of 5 independent experiments (n=12-16). *, p<0.05; **, p<0.01; *** p<0.001 (26RFa *vs* vehicle). $, p<0.05; $$, p<0.01; $$$ p<0.001 (insulin *vs* vehicle). #, p<0.05; ##, p<0.01(GPR103 antagonist *vs* vehicle).

The impact of i.c.v. administration of insulin and 26RFa on glycemia during a pyruvate tolerance test (IPPTT) was also investigated. Central injection of insulin induced a sustained inhibition of HGP that was not blocked by the co-administration of the 26RFa receptor antagonist 25e (Fig. 8A). In contrast to insulin, 26RFa did not alter the HGP, and the co-administration of the peptide with the GPR103 antagonist did not induce any change in the glycemic profile (Fig. 8B). Finally, we found that the inhibitory effect of central insulin on HGP was not altered in 26RFa-mutant mice as compared to their wild-type littermates (Fig. 8C).

**Figure 8.**
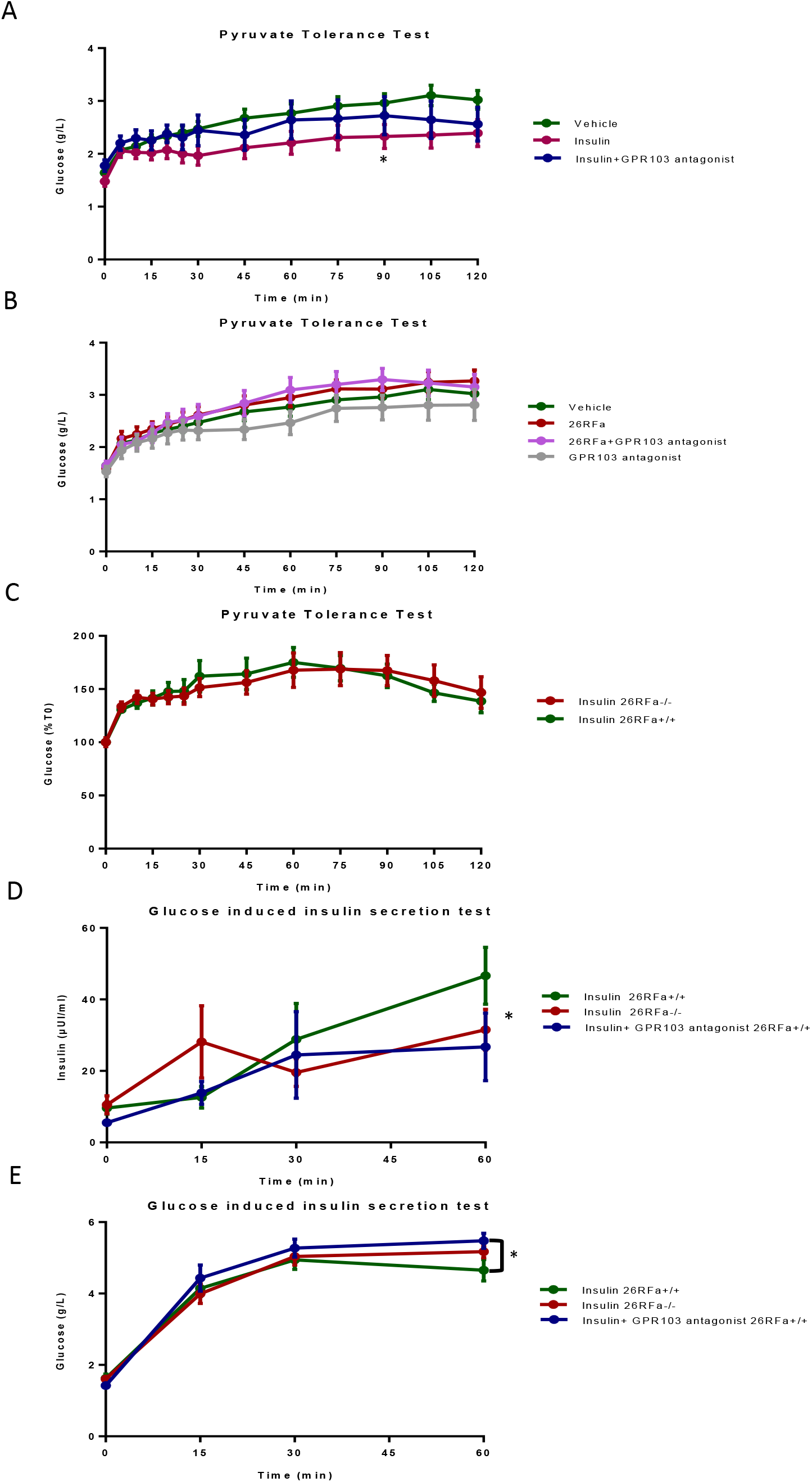
Central interaction between insulin and the hypothalamic 26RFa/GPR103 system during a pyruvate tolerance test and a glucose-induced insulin secretion test. **A:** Effect of i.c.v. administration of insulin (10 mUI) alone, or with the GPR103 receptor antagonist 25e (10^−4^ M) on plasma glucose levels during a pyruvate tolerance test (PTT). **B:** Effect of i.c.v. administration of 26RFa (3 µg) alone, or with the GPR103 receptor antagonist 25e (10^−4^ M) on plasma glucose levels during a PTT. **C:** Effect of i.c.v. administration of insulin (10 mUI) on plasma glucose levels during a PTT in 26RFa ^+/+^ and 26RFa^-/-^ mice. **D:** Effect of i.c.v. administration of insulin (10 mUI) alone, or with the GPR103 receptor antagonist 25e (10^−4^ M), on plasma insulin levels during a glucose induced-insulin secretion test (AGSIS) in 26RFa ^+/+^ and 26RFa^-/-^ mice. **E:** Effect of i.c.v. administration of insulin (10 mUI) alone, or with the GPR103 receptor antagonist 25e (10^−4^ M), on plasma glucose levels during an AGSIS in 26RFa ^+/+^ and 26RFa^-/-^ mice. Data represent means ± SEM of 5 independent experiments (n=9-15). *, p<0.05 (insulin *vs* vehicle or insulin vs insulin + GPR103 antagonist).

The effect of central insulin on its own peripheral production by the pancreatic islets was also examined (Fig. 8D). During an IPGTT, insulin injected centrally induced a significant increase (p<0.05) in peripheral insulin production that is blocked by the co-administration of the GPR103 antagonist (Fig. 8D). Similarly, insulin-induced insulin secretion was strongly attenuated in 26RFa-deficient mice (Fig. 8D). Measurement of plasma glucose levels during the same test indicated that glycemia displayed a mirror profile to that of insulinemia with more elevated plasma glucose levels in 26RFa^-/-^ mice and in animals co-injected with the 26RFa receptor antagonist (p<0.05), in comparison to wild type mice that received insulin centrally alone (Fig. 8E).

## Discussion

Although it is now recognized that the hypothalamus plays an important role in the global regulation of glucose metabolism in coordination with the pancreatic islets (1, 2), the molecular characterization of the neuronal circuits involved in the hypothalamic regulation of glucose homeostasis remains fragmentary. Indeed, accumulating evidence support the view that the AgRP and POMC neuronal systems of the Arc play a pivotal role in the central control of glucose metabolism (10, 11). However, physiological and genetic approaches indicate that these two peptidergic systems are not sufficient to explain and to understand the hypothalamic regulation of glucose homeostasis (11). In the present study, we provide the evidence that the 26RFa/GPR103 neuropeptidergic system of the hypothalamus is a novel actor of the central control of glucose homeostasis.

We firstly found that an i.c.v. administration of 26RFa induces a sustained antihyperglycemic effect that is associated with an increase of insulin production by the pancreatic islets without affecting insulin sensitivity. In addition, we show that this central antihyperglycemic effect of 26RFa is abolished when a GPR103 antagonist is co-administrated. Altogether, these findings indicate that 26RFa is involved in the brain regulation of glucose homeostasis, and that this central effect of 26RFa is mediated by the activation of its receptor GPR103. We previously showed that, at the periphery, 26RFa produced by the gut and the pancreatic islets displays a robust antihyperglycemic activity by acting as an incretin and by triggering insulin sensitivity (23-25). The present study suggests that the global antihyperglycemic effect of 26RFa is the result of a dual action of the peptide on the pancreatic islets and on the hypothalamus. 26RFa is not the single factor that is able to regulate glycemia by acting synergically at the peripheral and central level. For instance, it was reported that injection of glucose, leptin or insulin within the hypothalamus lowers blood glucose levels, increases liver insulin sensitivity (5,6,2), and that insulin decreases HGP (7). It was also observed that central injection of leptin (at doses too low to have any effect outside the brain) is able to normalize glycemia in mice with severe insulin deficiency (4). Hormones other than leptin can also act into the brain to promote insulin-independent glucose lowering. For instance, the gastrointestinal hormone FGF19 (or its rodent homologue, FGF15), when injected centrally, improves glucose tolerance in obese rats (30). Similarly, perfusion of the incretin GLP-1, in the Arc, causes a decrease of HGP (31). Recent studies also report an implication of the hypothalamic neuropeptide orexin in the brain regulation of glucose metabolism. Indeed, central administration of orexin A during the night-time awake period firstly elevates blood glucose levels and subsequently lowers daytime glycemia in normal and diabetic *db/db* mice via a bidirectional regulation of glucose hepatic production relayed by the autonomic nervous system (32, 33). Collectively, these observations support the new concept claiming that the control of glucose homeostasis results from a complex and highly coordinated interactions between the brain and the pancreatic islets.

26RFa has the interesting particularity to be produced at the periphery by the gut and the pancreatic islets (23-25), but also by a discrete neuronal population located in the hypothalamus (12, 16, 18, 19). This observation raises the possibility that 26RFa-producing neurons belong to the gluco-regulatory system of the hypothalamus and are responsible for the central effects of the peptide on glucose homeostasis. To test this hypothesis, we investigated whether peripheral factors well known to control the regulation of glucose metabolism by the hypothalamus may modulate the expression and release of 26RFa by hypothalamic neurons. For this, we developed two complementary approaches using the hypothalamic neuronal cell line m-HypoA-59 as a model of 26RFa-producing neurons and an *ex vivo* model of hypothalamic explants. Our experiments revealed that neither glucose, leptin nor insulin substantially impair 26RFa secretion by the m-HypoA-59 cells. Conversely, we found, using the *ex* and *in vivo* approaches that insulin drastically stimulates the expression and secretion of 26RFa by the hypothalamus. This finding strongly suggests that the 26RFa neuronal circuitry belongs to the gluco-regulatory system of the hypothalamus and may relay, at least in part, the central effects of insulin on the regulation of glucose metabolism. Consistent with this latter hypothesis, we show that 26RFa displays a very similar antihyperglycemic effect to that of insulin in terms of amplitude and kinetic, and that the central antihyperglycemic effect of insulin is abolished in presence of a 26RFa receptor antagonist and in mice deficient for 26RFa. In addition, we show that 26RFa-expressing neurons of the VMH and the LH also express the InsR and that central administration of insulin induces the expression of 26RFa in neurons of the VMH, LH but also of the Arc. Altogether, these complementary observations strongly suggest that 26RFa-expressing neurons are a target for insulin and mediate the regulatory effect of the hormone on glucose homeostasis within the hypothalamus. Interestingly, it was previously shown that injection of insulin into the Arc lowers blood glucose levels, increases liver insulin sensitivity (5, 6), and decreases HGP (7). The AgRP neurons of the Arc were identified as one of these neuronal populations targeted by insulin, as specific deletion of the insulin receptor in AgRP neurons partially blunts the ability of the hormone to inhibit HGP (11). By contrast, ablation of the insulin receptor in the POMC neurons of the Arc or specific activation of these neurons have no effect on glycemia nor HGP (11, 34). Here, we provide the evidence that insulin targets a new discrete population of hypothalamic neurons located in the VMH and the LH: the 26RFa-expressing neurons.

It is well documented that the central antihyperglycemic action of insulin is due, in part, to an inhibitory effect of the hormone on HGP (7). So, we examined whether the 26RFa neurons mediate the central insulin effects on HGP. We found that 26RFa injected centrally has no effect on HGP in contrast to insulin and that the effect of the hormone on HGP is not impaired by a blockade of the 26RFa receptor, nor in 26RFa-deficient mice. However, the literature reports that the central antihyperglycemic effect of insulin is also due to an activation of its own production by the pancreatic islets (35). In the present study, we confirm this statement and we show that GPR103 blockade or the absence of 26RFa (26RFa mutant mice) abolishes insulin-induced insulin production. These observations indicate that the hypothalamic 26RFa neurons are not involved in the central inhibitory effect of insulin on HGP but mediate the central effects of the hormone on its own peripheral production. The present findings, together with previous ones, raise the original hypothesis that insulin is able to target distinct neuronal populations in the hypothalamus, i.e. the AgRP neurons that mediate the inhibitory effect of the hormone on HGP, and the 26RFa neurons that mediate the stimulatory action of insulin on its own peripheral production. We propose that these two synergical actions may be responsible of the central antihyperglycemic effect of the hormone.

In conclusion, in the present study, we identified a novel hypothalamic neuronal circuitry that regulates glucose homeostasis, the 26RFa/GPR103 neuropeptidergic system and we provide evidence that this discrete neuronal population is a key relay for the central regulation of glucose metabolism by insulin.

## Methods

### Animals

26RFa^+/+^ and 26RFa^-/-^ male C57Bl/6J mice, weighing 22–25 g, were housed with free access to standard diet (U.A.R., Villemoisson-sur-Orge, France) and tap water. They were kept in a ventilated room at a temperature of 22±1°C under a 12-h light/12-h dark cycle (light on between 7 h and 19 h). All the experiments were carried out between 9 h and 18 h in testing rooms adjacent to the animal rooms. Mice were housed at three to five per cage. Unless otherwise stated, all tests were conducted with naïve cohorts of mice that were habituated to physiological and behaviour protocols before the beginning of experiments.

All experimental procedures were approved by the Normandy Regional Ethics Committee (Authorization: APAFIS#11752-2017100916177319) and were carried out in accordance with the European Committee Council Directive of November 24, 1986 (86/609/EEC).

### Animal surgery

Mice were anesthetized with isoflurane and placed in a stereotaxic frame. A stainless steel 26 gauge guide cannula (Phymep, Paris, France) was inserted into the right lateral ventricle (0.8 mm lateral to bregma and 2.2 mm ventral to dura mater). A stainless dummy small cap for 26-gauge cannula (Phymep) was inserted to prevent occlusion of the guide cannula and leakage of the cerebrospinal fluid. The cannula placement was secured with dental acrylic cement. At the end of the surgery, animals were treated with a nonsteroidal anti-inflammatory drug, buprenorphine after guide cannula placement. Following surgery, all animals were allowed to recover for two weeks, and were handled three times in order to be habituated to handling by researchers. Correct cannula positioning was confirmed by histological examination after Trypan blue injection. The injections were done using a Hamilton syringe (outer diameter 0.5 mm; Hamilton Co., Reno, NV) attached to polyethylene tubing. The i.c.v. injections were performed under anesthesia consisting of an i.p. administration of diazepam (5 mg/kg) and ketamine (100 mg/kg).

### Hypothalamic explants

The hypothalami, delineated by the posterior border of the optic chiasma, the anterior border of the mammillary bodies, and the fornix, were quickly dissected and removed. The hypothalamic explants were quickly removed and transferred into individual tubes containing 3 ml of artificial cerebrospinal fluid medium composed of 20 mM NaHCO_3_, 126 mM NaCl, 0.09 mM Na_2_HPO_4_, 6 mM KCl, 1.4 mM CaCl_2_, 0.09 mM MgSO_4_, 2 mM glucose, 0.18 mg/ml ascorbic acid, and 100g/ml aprotinin. The tubes were maintained at 37 C° in 95% O_2_, 5%CO_2_ under constant roughness. After a 4-h incubation with glucose (0.5 or 5 mM), leptin (10 or 100 nM) or insulin (10 or 100 nM), the culture media were collected and submitted to 26RFa assay, and the hypothalami stored at -80°C until qRT-PCR quantification.

### Cell culture

The [murine m-HypoA-59 neuronal cell line](https://www-sciencedirect-com.proxy.insermbiblio.inist.fr/topics/biochemistry-genetics-and-molecular-biology/murine) (CELLutions Biosystems Inc, Ontario, Canada) was used in the present study and cultured as previously described (27-29). For [mRNA](https://www-sciencedirect-com.proxy.insermbiblio.inist.fr/topics/medicine-and-dentistry/messenger-rna) quantification and 26RFa secretion studies, the cells were plated in 12-well plates (70×10^3^ cells/well and 200×10^3^ cells/well respectively). After 72 h, the cells were starved for 16 h in glucose-free DMEM (Sigma–Aldrich), supplemented with 2 mM glucose, 1% FBS, 1% penicillin/streptomycin, 3.7 g/L [sodium bicarbonate](https://www-sciencedirect-com.proxy.insermbiblio.inist.fr/topics/medicine-and-dentistry/sodium-bicarbonate), 0.584 g/L l-glutamate (Sigma–Aldrich) 0.11 g/L [sodium pyruvate](https://www-sciencedirect-com.proxy.insermbiblio.inist.fr/topics/medicine-and-dentistry/pyruvate-sodium). Test substances, i.e. glucose (0.5, 2 and 5 mM), leptin (0, 10 and 100 nM) and insulin (0, 10 and 100 nM), were added for 4 h in the culture medium. The supernatants were collected for 26RFa assay and the cells were subjected to RNA extraction for PCR quantification.

### Single-step Reverse Transcription PCR (RT-PCR) and quantitative RT-PCR (qRT-PCR)

Total RNA from hypothalamic explants of mice or m-HypoA-59 cells was isolated as previously described (25). Relative expression of the GPR103, 26RFa and insulin receptor (InsR) genes was quantified by real-time PCR with appropriate primers (Table 1). β-actin was used as internal control for normalization. PCR was carried out using Gene Expression Master Mix 2X assay (Applied Biosystems, Courtaboeuf, France) in an ABI Prism 7900 HT Fast Real-time PCR System (Applied Biosystems). The purity of the PCR products was assessed by dissociation curves. The amount of target cDNA was calculated by the comparative threshold (Ct) method and expressed by means of the 2-ΔΔCt method.

### Blood glucose and insulin measurements in mice

For measurements of basal glycemia and insulinemia, mice were fasted 6 h or 12 h before the test with free access to water. For i.p. glucose tolerance test, mice were fasted for 16 h with free access to water and then treated i.p. with glucose (2 g/kg). For insulin tolerance test, mice were fasted for 6 h before the test with free access to water, and then injected i.p. with 0.75 units/kg body weight of human insulin (Eli Lilly, Neuilly-sur-Seine, France). For pyruvate tolerance test, mice were fasted for 16 h before the test with free access to water and then injected i.p. with sodium pyruvate (2 g/kg; Sigma Aldrich). Test substances including 26RFa (3 µg), 25e (10^−4^ M) and insulin (10 mUI) were dissolved in Hepes buffer and injected i.c.v. in a 2 µl volume. Plasma glucose concentrations were measured from tail vein samplings at various times using an AccuChek Performa glucometer (Roche Diagnostic, Saint-Egreve, France). Plasma insulin concentrations (from tail vein samplings) were determined using an ultrasensitive mouse insulin AlphaLisa detection kit (cat number AL204C) from Perkin Elmer.

### 26RFa radioimmunoassay

Quantification of 26RFa in culture media was carried out using a specific radioimmunoassay (RIA) set up in the laboratory (3). Briefly, each culture medium sample was diluted (1:1) in a solution of water/TFA (99.9:0.1; v/v) and pumped at a flow rate of 1.5 ml/min through one Sep-Pak C_18_ cartridge. Bound material was eluted with acetonitrile/water/TFA (50:49.9:0.1; v/v/v) and acetonitrile was evaporated under reduced pressure. Finally, the dried extracts were resuspended in PBS 0.1 M and assayed for 26RFa.

### RNAscope-In situ hybridization experiments

26RFa^+/+^ or ^-/-^ mice were deeply anesthetized and perfused transcardially with sterile Phosphate Buffer Saline followed by ice-cold phosphate buffered 4% paraformaldehyde (PFA, pH 7.4) 45 mins after i.c.v. administration of hepes buffer or insulin (10 mU). The brains were removed, post-fixed overnight at 4°C in sterile 4% PFA, transferred to sterile 15% sucrose (12 h) and then to 30% sucrose (12 h) in 0.1 M PBS (pH 7.4) at 4°C. 12 µm-thick sections were cut on a cryostat and mounted on SuperFrostPlus slides and subsequently stored at -80°C. Fluorescent in situ hybridization for the simultaneous detection of the 26RFa, the InsR and AgRP transcripts was performed using RNAscope (ACD; Advanced Cell Diagnostics). The InsR probe targeted the region 7059-8053 (accession number NM_010568.2; ACD, Cat No. 401011-C2). The 26RFa (Qrfp) probe targeted the region 112–1081 (accession number NM_183424.4; ACD, Cat No. 464341-C3). The AgRP probe targeted the region 11–764 (accession number NM_001271806.1; ACD, Cat No. 400711). Negative and positive control probes recognizing dihydrodipicolinate reductase, DapB (a bacterial transcript) and PolR2A (C1 channel), PPIB (C2 channel) and UBC (C3 channel), respectively, were processed in parallel with the target probes to ensure tissue RNA integrity and optimal assay performance. In addition, the specificity of the 26RFa probe was assessed by performing an hybridization of brain sections of 26RFa^-/-^ mice that resulted in an absence of 26RFa mRNA signal (Fig. S1). All pretreatment reagents, detection kit, and wash buffer were purchased from ACD. All incubation steps were performed at 40°C using a humidified chamber and a HybEz oven (ACD). The day of the assay, all the hybridization steps, i.e., target retrieval, dehydration, probe hybridization, amplification steps, and detection of the probe, were performed according to the online protocol for RNAscope Multifluorescent Assay. Briefly, the procedure included the following steps: slides were first submerged in Target Retrieval (ACD) heated to 98.5°C– 99.5°C for 5 min, followed by two brief rinses in sterile water. The slides were quickly dehydrated in 100% ethanol and allowed to air dry for 5 min. The next day, the sections were incubated with Protease III for 30 min (ACD). The AgRP probe (channel 1), the InsR probe (channel 2) and the 26RFa probe (channel 3) were mixed according to the manufacturer’s instructions and hybridized to the sections for 2 h, followed by 2 min washes in wash buffer (ACD), incubation with Amp1 for 30 min, two washes, Amp2 for 30 min, two washes and finally Amp3 for 15 min followed by two washes. Sections were then incubated with amplification system (HRP and Opal fluorophore). The AgRP probe was detected with Opal 520 (Akoya Biosciences, FP1487001KT), the InsR with Opal 650 (Akoya Biosciences, FP1496001KT) and the 26RFa probe with Opal 570 (Akoya Biosciences, FP1488001KT). Sections were then immediately cover slipped with Fluoromount Aqueous Mounting (Sigma, F4680) with DAPI and stored in dark at 4°C until imaging.

Images were captured using a confocal Leica TCS SP-8-X microscope equipped with a 40x objective and a Leica Thunder Imager 3D (DM6 B). Maximum intensity projections were made inFIJI (NIH), and images were adjusted for brightness and contrast.

### Statistical analysis

Statistical analysis was performed with GraphPad Prism (version 6). A Kruskal Wallis test with Dunn’s multiple comparison test or Ordinary one-way ANOVA with Sidak’s multiple comparison were used for comparisons between different groups. ANOVA two-ways were used for repeated measures. A post-hoc comparison using a Bonferoni, Tukey or Sidak test was applied according to ANOVA results. All data represent means ± SEM. Statistical significance was set up at p < 0.05.

## Data and resource availability

### 1/ Data sharing

All data generated or analyzed during this study are included in the published article (and its online supplementary files).

### 2/ Resource sharing

No applicable resources were generated or analyzed during the current study.

## Author Contributions

M.E.M., M.P. and N.C. contributed to the study design and interpretation, and wrote the manuscript. S.T., J.M., A.A., M.P., MA. LS. and M.E.M. performed the *in vivo* experiments on mice. S.T., C.D., M. D. and M.P. contributed to the RNAscope experiments. M. D. and M.E.M. contributed to the PCR experiments. A. N. provided the GPR103 antagonist. A. B. and E. N. performed the insulin assays, and J. L. and B.L. produced synthetic 26RFa. G.P., A. N. and Y. A. revised and approved the final version of the manuscript. N.C. is the guarantor of this work and, as such, had full access to all the data in the study and takes responsibility for the integrity of the data and the accuracy of the data analysis.

