## Supplemental Fig.1 for "Identification of a discrete neuronal circuit that relays insulin signaling into the brain to regulate glucose homeostasis"

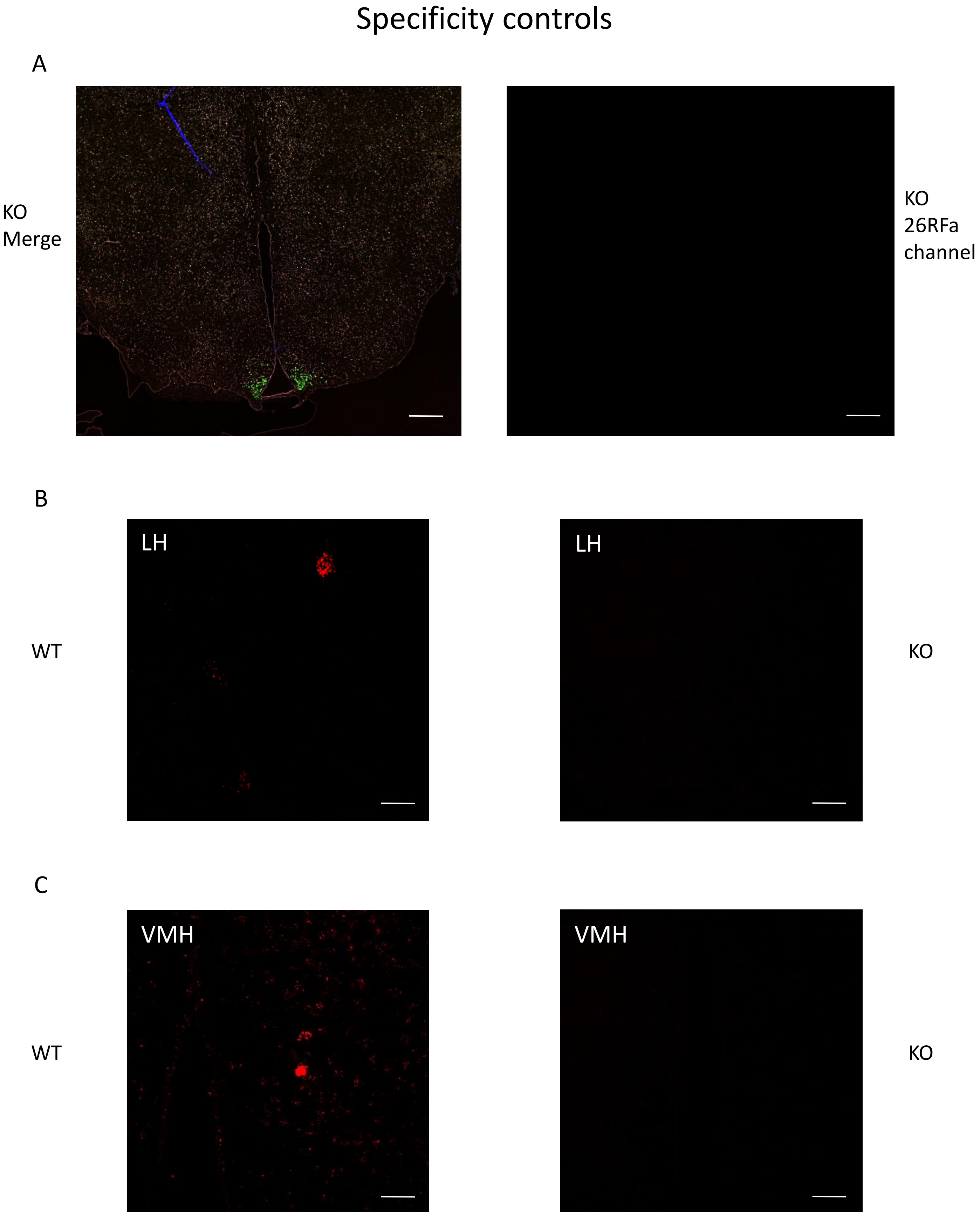


**Figure S1. Specificity controls of the 26RFa mRNA probes used during the RNAscope procedure. A:** Hypothalamic section of a 26RFa$^{-/-}$ mouse (KO) treated with 26RFa (red), AgRP (green) and insulin receptor (InsR) (white) probes for the RNAscope approach showing the total absence of red labelling. **B, C:** Comparison of the 26RFa mRNA labelling (red spots) between wild type (WT) and 26RFa $^{-/-}$ (KO) mice in the lateral hypothalamic area (LH) and the ventromedial hypothalamic nucleus (VMH) showing no labelling in the brains of the 26RFa mutant mice. Scale bars: (A)=200 µm; (B, C)=25 µm.
